# Proprotein Convertase Subtilisin/kexin Type 9 Links Inflammation to Vascular Endothelial Cell Dysfunction

**DOI:** 10.1101/2021.01.15.426820

**Authors:** Thorsten M. Leucker, Nuria Amat-Codina, Stephen Chelko, Gary Gerstenblith

**Author notes:** **Correspondence to:** Thorsten M. Leucker, M.D. Ph.D., Blalock 547, Johns Hopkins Hospital, Division of Cardiology, 600 N. Wolfe Street, Baltimore, MD 21287. Dr. Chelko’s current address is: Department of Biomedical Sciences, Florida State University College of Medicine, Tallahassee, FL, 32306.

## Abstract

Vascular endothelial cell (EC) dysfunction is a pathological mediator of he development, progression, and clinical manifestations of atherosclerotic disease. Inflammation is associated with EC dysfunction, but the responsible mechanisms are not well characterized. There is substantial evidence that serum proprotein convertase subtilisin/kexin type 9 (PCSK9) is increased in pro-inflammatory states and that elevated PCSK9 levels are associated with adverse cardiovascular outcomes after controlling for traditional risk factors, including low-density lipoprotein (LDL) cholesterol.

Here we investigate PCSK9 as a novel link between inflammation and vascular EC dysfunction, as assessed by nitric oxide (NO) bioavailability. Tumor necrosis factor alpha (TNF-α), a pro-inflammatory cytokine, increased *PCSK9* mRNA expression and PCSK9 protein levels in isolated human aortic ECs, which were accompanied by reduced total and phosphorylated endothelial nitric oxide synthase (eNOS) protein levels and NO bioavailability. Finally, genetic PCSK9 reduction utilizing a PCSK9 specific siRNA in human aortic ECs resulted in the rescue of phosphorylated eNOS protein levels and NO bioavailability.

Our results demonstrate that PCSK9 is increased in human aortic ECs exposed to a pro-inflammatory stimulus and that this increase is associated with EC dysfunction. Silencing of TNFα-mediated augmentation of *PCSK9* expression utilizing a small interfering RNA against *PCSK9* rescued the inflammation-induced EC dysfunction. These results indicate that PCSK9 is a causal link between inflammation and EC dysfunction, a potent driver of atherosclerotic cardiovascular disease.

## Introduction

Atherosclerotic cardiovascular disease (ASCVD) is a leading cause of premature mortality and significant lifelong disability in the United States. [1] Clinical and experimental data support a critical role for inflammation in the development and progression of ASCVD. [2] Vascular endothelial cell (EC) dysfunction is an early manifestation of atherosclerosis [3] and inflammation is associated with EC dysfunction. [4] Identifying novel, safe, and intervenable links between pro-inflammatory conditions and vascular EC dysfunction is an unmet need.

Proprotein convertase subtilisin/kexin type 9 (PCSK9) is a serine protease originally discovered as neural apoptosis-regulated convertase-1 (NARC-1) during embryonic brain development. [5] Several subsequent studies defined the important role of PCSK9 in cholesterol metabolism, [6] which is mediated by its ability to bind to the epidermal growth factor precursor homology domain-A (EGF-A domain) of the low-density lipoprotein (LDL)-receptor (LDLR), targeting it for lysosomal-mediated degradation. [7] Leander et al. reported that serum PCSK9 levels are associated with the risk of future cardiovascular events independent of established cardiovascular risk factors, including serum LDL-cholesterol [8]. Serum PCSK9 levels are elevated in various inflammatory conditions [9] and *PCSK9* is expressed in vascular EC. [10, 11]

We tested the hypothesis that vascular EC PCSK9 links inflammation with vascular EC dysfunction.

## Methods

### Human Study Approval

The following human samples were obtained during coronary artery bypass surgery: left internal mammary artery (LIMA) from individuals with established atherosclerotic cardiovascular disease. The study was approved by the Johns Hopkins School of Medicine institutional review board (IRB Study No. NA_IRB00164620 to TL and GG). All participants provided written informed consent.

### Cell culture

Human aortic endothelial cells (HAEC) were obtained from Cell Applications, Inc San Diego, CA and maintained in Human Endothelial Cell Medium culture medium (cat#, 213-500 Cell Applications, Inc.) according to the manufacturer’s protocol. Lipofectamine RNAiMAX reagent (ThermoFisher Scientific, cat# 13778030) was used for transfection of HAECs with *PCSK9* siRNA according to the manufacturer’s protocol described below.

### Western immunoblotting

HAEC cells (passage 3-4) were lysed in ice-cold RIPA buffer (ThermoFisher, cat# 89900) containing protease and phosphatase inhibitors (ThermoFisher, A32959, Waltham, MA). Protein lysates were incubated with 1X NuPage LDS sample buffer (ThermoFisher, cat# NP0008) containing 1X NuPage reducing agent (ThermoFisher, cat# NP0004), boiled for 15 min at 95C prior, then proteins were separated on a NuPAGE 4-12% Bis-Tris gel (ThermoFisher, cat# NP0321). Gels were transferred onto PVDF membranes and the following primary antibodies were used: Endothelial NO synthase (eNOS, ab76198, 1:500 dilution, Abcam), phosphorylated eNOS (p-eNOS at Serine [S1177], ab184154, 1:500 dilution, Abcam), PCSK9 (ab28770, 0.3ug/ml dilution, Abcam), VE-Cadherin (ab33168, 1:1000, Abcam), β-actin (ab8226, 1:5,000 dilution, Abcam), GAPDH (D16H11, 1:2,000 dilution, Cell Signaling Technology).PVDF membranes were incubated overnight at 4C with primary antibodies diluted in 1X TBS Intercept Blocking Buffer (LiCor, cat# 927-60001). The following day, PVDF membranes were washed three times in 1X TBST, then incubated with species-specific IRDye secondary antibodies for 1hr at room temperature (LI-COR, at 1:5,000).

### Endothelial Cell Dysfunction Assay

Nitric oxide (NO) bioavailability was measured in HAEC culture medium to assess the extent of EC dysfunction, as described previously. [12] Specifically, the measurement of combined nitrite and nitrate levels were used as the amount of NO bioavailability using the Nitric Oxide Assay kit (cat# EMSNO, ThermoFisher).

### RT-PCR and real-time PCR

TRIzol (Thermo Fisher Scientific) was used to extract total RNA from HAEC. RNA was reverse transcribed using the High-Capacity cDNA Reverse Transcription Kit (ThermoFisher, ABI 4368814). *PCSK9* mRNA expression was assessed by real-time PCR using *PCSK9* primer sets 5’-TGTCTTTGCCCAGAGCATC-3’ and 5’-GTCACTCTGTATGCTGGTGTC-3 (Integrated DNA Technologies) and SYBR Select Master Mix (ThermoFisher, ABI 4472908).

### PCSK9 Knockdown

The DsiRNA used was obtained from IDT (Coralville, Iowa, USA). Transfection efficiency was obtained by transfecting HPRTsiRNA, non-silencing siRNA and TYR-dye into ECs for a 24 hrs. HAEC were plated in 6-well plates and maintained until a 80%-90% confluency was achieved. Cells were then treated with either 10nM of non-silencing control siRNA or 10 nM *PCSK9* siRNA using Lipofectamine RNAiMAX reagent (ThermoFisher, cat# 13778030). Cells were incubated at 37C, 5% CO2 for 48 hours. Following control or *PCSK9* siRNA incubation, cells were treated with 20 ng of TNF-α (R&D systems, 10291-TA-020) for 24 hours. Cells and cell culture medium were then collected for downstream proteomic and biomolecular assays.

### Immunofluorescence

Vascular tissue was fixed in 4% paraformaldehyde and embedded in paraffin blocks. Samples were sectioned at 5μm, deparaffinized using Xylene, rehydrated (95%, 70%, and 35% ethanol), 1X PBS wash, antigen retrieval, blocked for 1 hr at room temperature in 1X PBS containing 0.1% Triton-X and 5% BSA (blocking buffer), then incubated with 5ug/ml VE-Cadherin and PCSK9 primary antibodies at 4C overnight, as previously described.[13] The following day, slides were washed three times in 1X PBS, and incubated with species-specific Alexa Fluor 647 and 488 secondary antibodies at room temperature for 1 hr. Slides were washed three times with 1X PBS then Fluoroshield with DAPI mounting medium (Sigma, F6057) was used to stain for nuclei and affix coverslips, respectively. Immunofluorescence images were obtained using a Leica TCS SPE II laser scanning confocal microscopy.

### Statistics

All statistical analyses were performed with GraphPad Prism 9 software. Data were tested for normality using the Shapiro-Wilk test. Parametric (Student t-test) and nonparametric (Kolmogorov-Smirnov test) tests were used when appropriate for normally distributed and skewed data, respectively. Normally distributed data are presented as mean±StDev and skewed data are presented as medians and interquartile ranges. Statistical significance was defined as P≤0.05.

## Results

### PCSK9 is abundant in vascular tissue of patients with atherosclerotic cardiovascular disease

Immunohistochemistry staining of left internal mammary artery (LIMA) samples discarded during coronary artery bypass surgery demonstrated vascular distribution of PCSK9. Figure 1 shows representative images of a LIMA obtained from a 67-year-old man with atherosclerotic cardiovascular disease. Panel A and zoomed image panel B show cross sectional views of the artery with PCSK9 immuno-staining (green) along the inner and outer elastic membranes. Panels C and D (zoomed images) depict co-localization of PCSK9 (green) and vascular endothelial cadherin (purple). The images also demonstrate additional localization of PCSK9 in the tunica adventitia.

**Figure 1.**
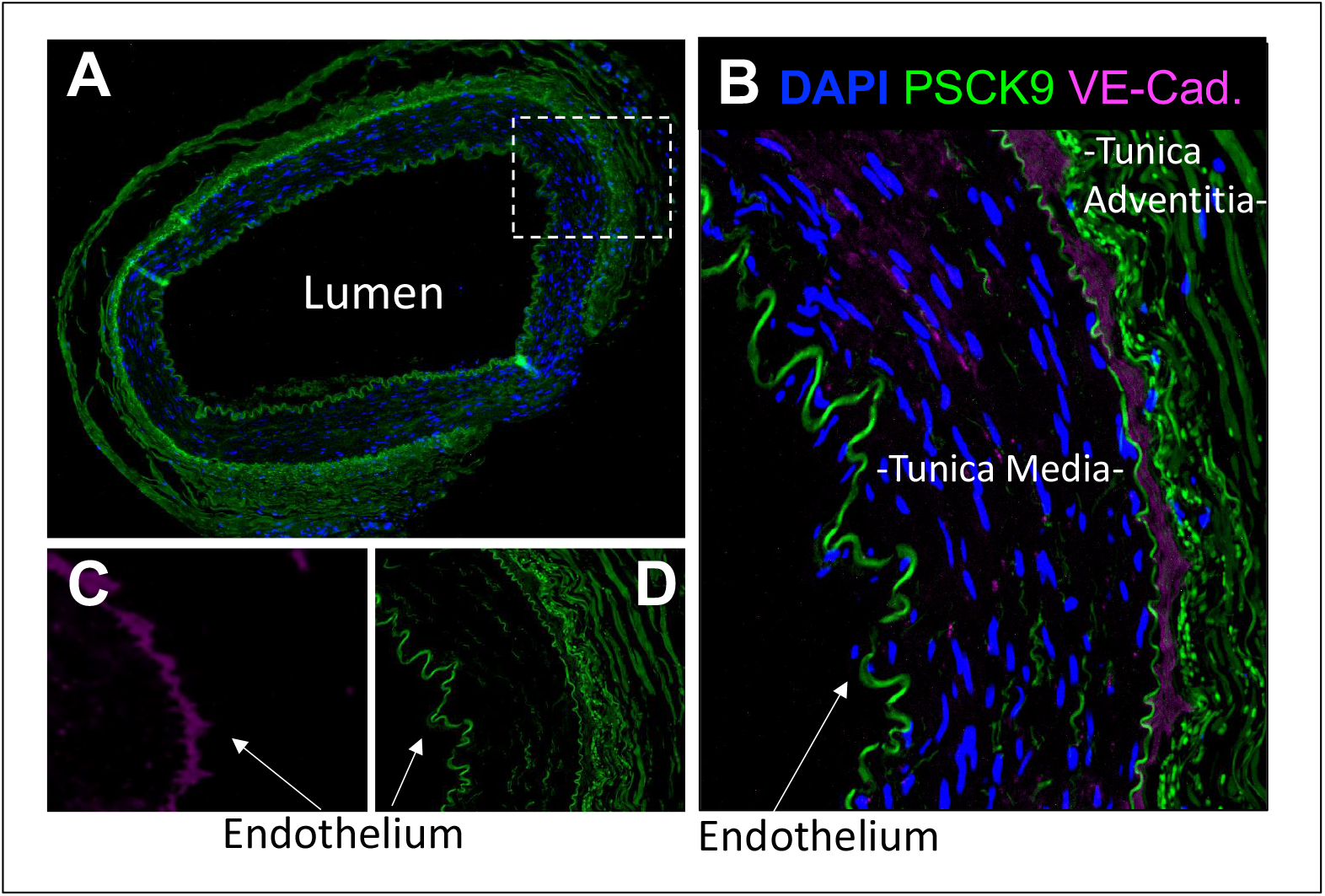
PCSK9 is present in vascular endothelium. Depicted images are of left internal mammary artery (LIMA) segments from a 67-year-old man with atherosclerotic cardiovascular disease undergoing coronary artery bypass grafting. **A**. and zoomed image **B.** representative confocal immunofluorescence images from the LIMA demonstrating PCSK9 immuno-staining (green) along the inner and outer elastic membranes, Zoomed images of LIMA **C and D** demonstrating co-localization of PCSK9 (green) and vascular endothelial cadherin (purple); DAPI= 4′,6-diamidino-2-phenylindole; PCSK9= Proprotein convertase subtilisin/kexin type 9; VE-Cad.= vascular endothelial cadherin).

### Endothelial cell PCSK9 mRNA is increased after exposure to pro-inflammatory stimulus

ECs obtained from individuals 56-69 years old were used for the experiments shown in Figure 2 (n= 3 separate EC lines with 2-3 replicates per experiment). Panel A demonstrates that TNF-α stimulates EC PCSK9 mRNA production (1.98 [0.74, 3.36]-fold increase p=0.02 vs. control). Furthermore, pre-treatment of ECs with PCSK9 directed siRNA shows a 71.07% [15.08, 30.16] reduction in baseline PCSK9 mRNA expression (p<0.01 vs. scrambled siRNA control) and no TNF-α induced increase.

**Figure 2.**
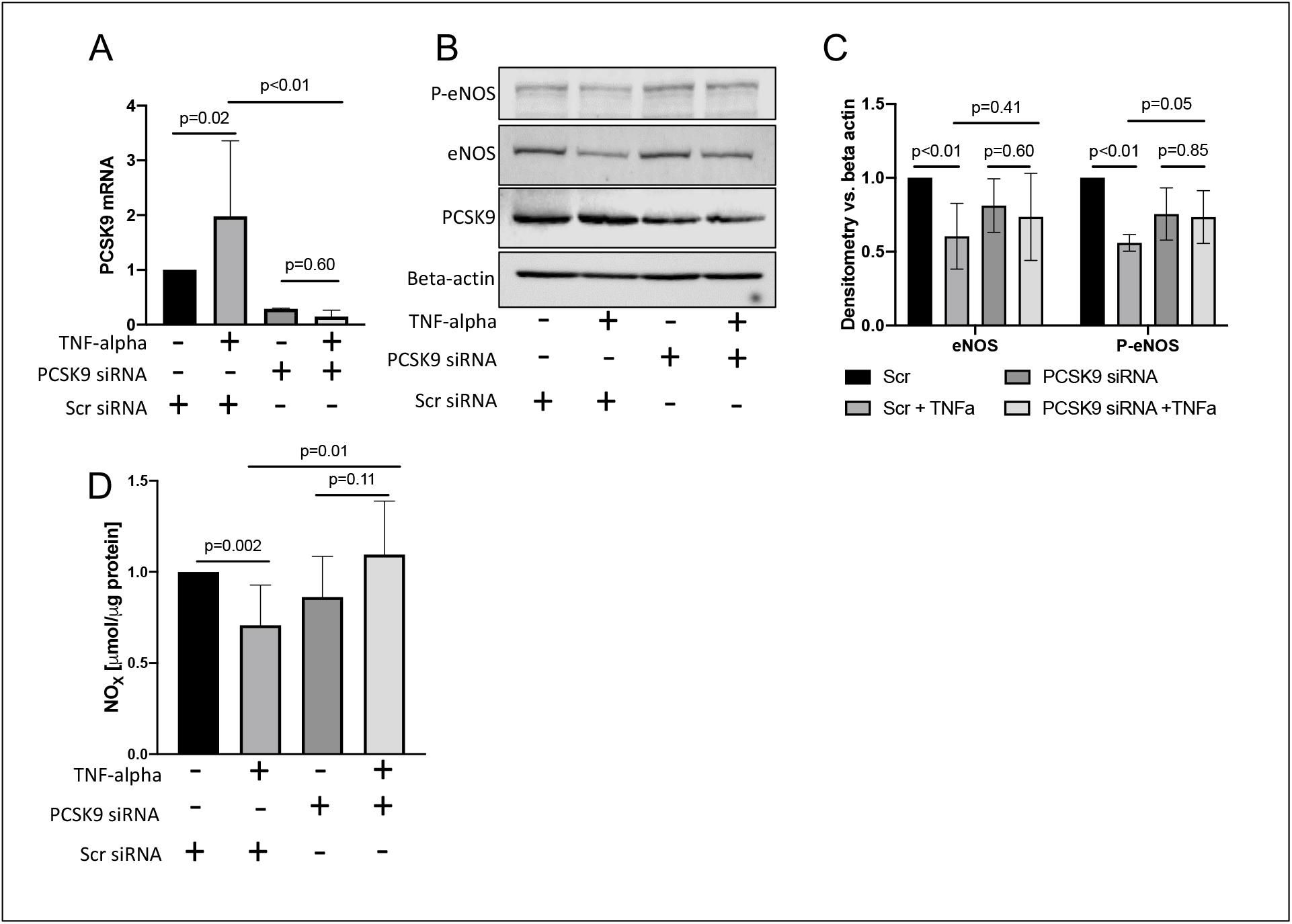
EC PCSK9 knock down rescues inflammatory induced EC dysfunction. Human aortic ECs were exposed to the pro-inflammatory stimulus, TNFa (20ng/mL), in the presence and absence of genetic PCSK9 knock-down (PCSK9 siRNA). **A**. PCSK9 mRNA levels were determined by real-time PCR; **B.+ C**. Immunoblot with phospho eNOS (P-eNOS), total eNOS, PCSK9 and beta actin antibodies; **D.** EC cell culture media was collected for nitrite and nitrate measurements; Data are presented as median ± IQR (Panel A) and mean ± SD (Panel C and D); Scr = scrambled; N=3 using ECs from 3 different donors with 2-3 replicates per experiment.

### Genetic EC PCSK9 interference rescues TNF-α induced EC dysfunction

Figure 2 B shows representative Western blot images of ECs treated with TNF-α in the presence of PCSK9 small interfering RNA (PCSK9 siRNA) and in its absence (scrambled siRNA). TNF-α reduces EC nitric oxide synthase (eNOS) to 60.47% ± 22.26% of baseline, p<0.01 vs. control, and Akt-mediated phosphorylated eNOS-Ser 1177 to 55.91% ± 5.62% of baseline, p<0.01 vs. control, in the presence of scr siRNA. However, there is no significant decrease in total eNOS (p=0.60) or phosphorylated eNOS in the presence of PCSK9 siRNA (p=0.85; Fig.2 C). Panel D shows that TNF-α reduces EC nitric oxide (NO) production by 29.25% ± 22.1% compared to control, p=0.002. However, NO bioavailability is not decreased in the presence of genetic PCSK9 silencing.

## Discussion

We demonstrate, in human vascular ECs, that TNF-α, a typical pro-inflammatory stimulus, induces vascular EC PCSK9 expression leading to reduced eNOS phosphorylation and EC NO bioavailability. The adverse pro-inflammatory effects on EC function are not present following genetic PCSK9 silencing. As the experiments were performed in isolated cells with normal media, the findings are independent of the known effects of PCSK9 on cholesterol metabolism.

Inflammation is a well-recognized important cardiovascular risk factor, [2] which is likely mediated, in part, by impaired EC-derived NO bioavailability. [4] NO exerts multiple direct anti-atherosclerotic effects including inhibition of LDL-oxidation [14] and of inflammatory cell adhesion to the arterial endothelial surface [15] and migration of the cells into the vascular wall. [16] Nitric oxide also inhibits platelet activation and thrombus formation. [17, 18]

Although inflammation is known to impair EC function, [4] the responsible mechanisms are not well characterized. Serum PCSK9 is increased in patients with pro-inflammatory states [9] and, in a murine model, inflammatory stimuli raised hepatic PCSK9. [19] In an observational study, serum PCSK9 levels in those with stable coronary disease was associated with future cardiovascular events [20] and in a prior clinical study, we demonstrated that in people with HIV, a pro-inflammatory condition, short term treatment with a monoclonal antibody against PCSK9 significantly improved NO-dependent coronary artery EC function. [21]

Our results provide further mechanistic insight regarding the impact of inflammation on EC function and the role of PCSK9 in mediating that impact. To our knowledge, this is the first report demonstrating that PCSK9 links inflammation to vascular EC dysfunction.

